# Early-life stress-induced gut dysbiosis is ameliorated by nicotinamide treatment

**DOI:** 10.64898/2026.08.19.745716

**Authors:** Stuti Srivastav, Pratik R. Chaudhari, Shital Suryavanshi, Nishtha Pange, Vidita A. Vaidya, Amitesh Anand

## Abstract

Early life stress (ELS) in the form of adverse experiences in childhood results in multiple psycho-physiological pathologies during adulthood and accelerates aging. Such pathologies are recently observed to be alleviated upon nicotinamide treatment in a maternal separation model of ELS. We report a nicotinamide-driven amelioration of gut dysbiosis in middle-aged rodents with a history of neonatal maternal separation.

## Main

With advancing age, human development largely shifts from the formative to the regenerative phase; thus, early life represents a critical period of growth ^1,2^. This developmental window lays the foundation for an individual’s physical, physiological, and cognitive traits, with perturbations during this period often resulting in lifelong consequences ^3,4^. Stressful experiences in this critical phase, in the form of neglect, abuse, infection, etc., significantly increase the lifetime risk for psychopathology. Between 2010 and 2023, 400 million children under the age of five suffered from physical and/or psychological violence globally ^5^. Early-life stress (ELS) is linked to increased incidence of psychiatric disorders, substance abuse, irritable bowel syndrome, and metabolic syndrome in adulthood ^6–10^. Thus, it becomes pertinent to examine potential therapeutic strategies targeting early stress-related pathologies in adulthood.

Animals with a history of ELS exhibit an increase in systemic inflammation, mitochondrial dysfunction, and blood-brain-barrier permeability. Such animals are reported to show cognitive decline, neurodegeneration, and signatures of premature aging ^11,12^. We have recently observed that supplementation with nicotinamide, which is a mitochondrial potentiator, rescues the inflammatory and premature-aging-related pathologies of ELS in rodents ^16^. Notably, nicotinamide adenine dinucleotide (NAD) level decreases with age and recovery of NAD concentration aids in mitigating senescence-associated disorders ^17,18^. Nicotinamide utilisation is intricately linked to the gut microbiome, as several enzymes of microbial origin, such as nicotinamidase (*pncA*), are critical for NAD metabolism ^19^. The gut microbiota is necessary for the manifestation of neurocognitive symptoms associated with ELS in mice and is central to both stress-related inflammation and aging ^13–15^. Therefore, we were motivated to examine the involvement of the gut microbiota in alleviating ELS pathologies upon nicotinamide supplementation.

The rodent-based maternal separation (MS) model recapitulates human ELS morbidities ^20^. During MS, rat pups are separated from their dams for three hours daily from postnatal day 2 till 14, and pathophysiological assessment is performed later in their life (Fig. 1A). Given that the pathologies of ELS manifest well beyond young adulthood, we maintained maternally separated rats into middle aged life ^16,21^. We examined the changes in the gut microbiome by performing 16S rRNA amplicon sequencing of fecal samples at two epochs of life: young adulthood and middle age (Fig 1A). We compared microbial diversity measures across the age groups and treatment paradigms using alpha and beta diversity indices.

**Figure 1.**
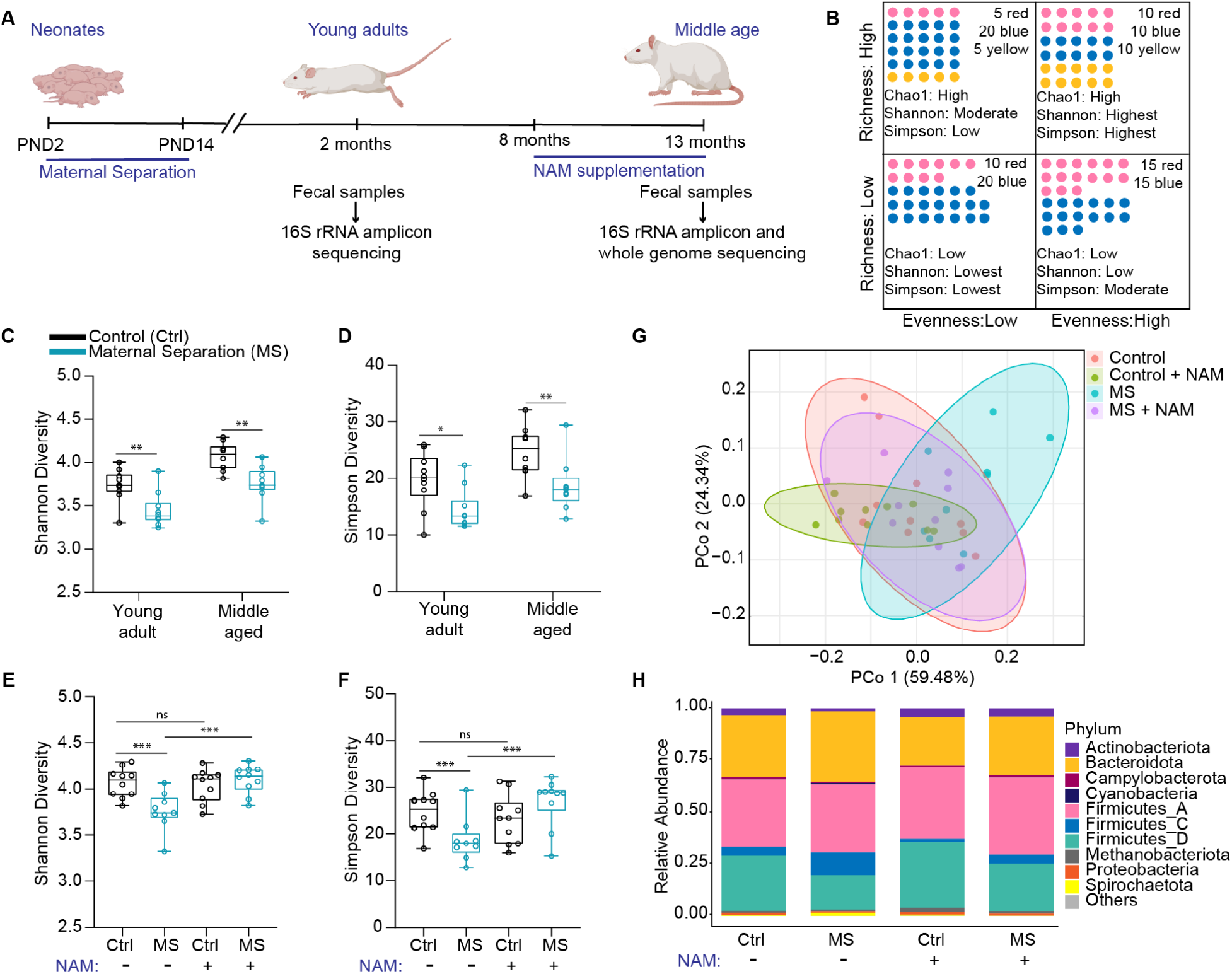
Maternal separation-induced gut dysbiosis signatures are reversed with nicotinamide supplementation. (A) Schematic depicting study design. Male Sprague Dawley rats were subjected to neonatal maternal separation (MS) from postnatal day (PND) 2 to 14. Fecal samples from these rats were collected at young adulthood (2 months old) and middle age (13 months old) for performing metagenomics analyses as indicated. Nicotinamide (NAM) was supplemented in drinking water from 8 to 13 months of age. (n = 9-10 for all groups). (B) Schematic representation of alpha diversity indices used in this study. Circles of various colours represent different taxa. (C&D) Box plots representing changes in the Shannon or Simpson diversity indices in gut microbiomes from control and MS animals at young adulthood and middle aged timepoints. (E&F) Box plots representing changes in the Shannon or Simpson diversity indices in gut microbiomes from control and MS animals at middle age with NAM supplementation. (C-F: n=9-10 biological replicates, box plots extend from 25^th^ to 75^th^ percentiles; whiskers represent minimum and maximum data points. Statistics were performed using the Wilcoxon rank-sum test with Bonferroni post-hoc correction. ns: *p*-value > 0.05, *: *p*-value ≤ 0.05, **: *p*-value ≤ 0.01, and ***: *p*-value ≤ 0.001). (G) Bray-Curtis principal coordinate analysis (PCoA) plot for control and MS animals at middle-age with and without NAM supplementation. (H) Stacked bar plots representing the relative abundances of the top 10 most abundant phyla across the groups mentioned on the x-axis.

We assessed alpha diversity measures of the gut microbiome using the Chao1, Shannon, and Simpson diversity indices (Fig. 1B). The Chao1 index represents taxa richness while the Shannon and Simpson indices also take into account the evenness of taxa. While the Shannon diversity is more sensitive to richness over evenness, the Simpson diversity depends more on taxa evenness compared to richness (Fig. 1B)^22^. As expected, an age-wise progression in various indices of alpha diversity was observed in both control and MS groups (Fig 1C-D, Suppl. Fig. 1A). However, across young adulthood and middle age, animals with a history of MS had an overall lower gut microbiome alpha diversity compared to the respective age-matched controls (Fig. 1C-D; Suppl. Fig. 1A). Encouragingly, these observations corroborate previous studies showing that neonatal MS causes a decreased gut microbial diversity in young adult rodents ^23^. We further establish that this dysbiosis state is persistent till middle age. This alteration in gut microbial diversity appears to precede the onset of inflammatory, metabolic, and neurocognitive defects observed after MS during middle age ^16^.

Next, we examined the influence of *ad libitum* oral nicotinamide supplementation on the gut microbiome of middle-aged control and MS rats (Fig. 1A). Nicotinamide supplementation did not seem to affect the Chao1 index; however, there was a significant trend towards restoration of the Shannon and Simpson diversities (Fig. 1F and G; Suppl. Fig. 1B).

In beta diversity, which compares microbial diversity between samples, the Jaccard index of various treatment groups largely overlapped with each other (Suppl. Fig. 1C). This overlap is expected as the Jaccard index does not factor taxa abundance, the primary metric noted to be influenced by nicotinamide treatment. We then performed Bray-Curtis principal coordinate analysis (PCoA), which accounts for taxa abundance also. The MS group was most segregated along the first principal coordinate compared to the control group and the NAM-supplemented MS group (Fig. 1G). The overlap of control and NAM-supplemented MS groups suggests a likely shift of the microbial abundances to a healthy control-like state on nicotinamide supplementation.

We then narrowed our analysis to phylum-level taxonomic abundance assessment. The relative abundance of various phyla was affected by maternal separation and nicotinamide supplementation (Fig. 1H). We examined the distribution profile of the top ten most abundant phyla that were common across groups. Maternal separation was observed to reduce the relative abundances of Actinobacteriota, Firmicutes_D, and Proteobacteria when compared to healthy controls. Whereas the phyla Bacteroidota, Firmicutes_C, and Spirochaetota showed increased relative abundance after maternal separation. Strikingly, nicotinamide supplementation altered the relative abundance of many phyla that were influenced by MS (Fig. 1H). Specifically, in animals with a history of MS, nicotinamide supplementation increased the relative abundances of Actinobacteriota and Firmicutes_D while reducing the abundances of Bacteroidota, Firmicutes_C, and Spirochaetota. Overall, the treatment with nicotinamide appears to reverse the changes in taxa abundance caused by MS. Interestingly, some of these alterations, like the reduction in Bacteroidota and increase in Firmicutes_D, were noted in control animals also, suggesting that nicotinamide supplementation favours a specific shift in bacterial abundances.

To resolve the impact of nicotinamide supplementation at lower taxonomic levels, we performed shotgun whole genome sequencing using fecal samples from middle-aged animals (Fig. 1A). At species level, the relative abundance of *Segatella copri* (previously *Prevotella copri*) increased in the microbiomes of rodents with a history of MS (Fig. 2A). Meanwhile, the relative abundances of *Bifidobacterium pseudolongum, Ligilactobacillus faecis*, and *L. murinis* decreased in the gut microbiome after MS. Consistent with the phylum-level observations, adulthood nicotinamide supplementation reversed the abundance changes in several of these species (Fig. 2A). Specifically, nicotinamide supplementation decreased the relative abundance of *S. copri* while increasing *B. pseudolongum, L. faecis*, and *L. murinis* abundance. Notably, abundance of Bifidobacteria and Lactobacilli is observed to decrease with aging in humans and rodents ^15,24^.

**Figure 2.**
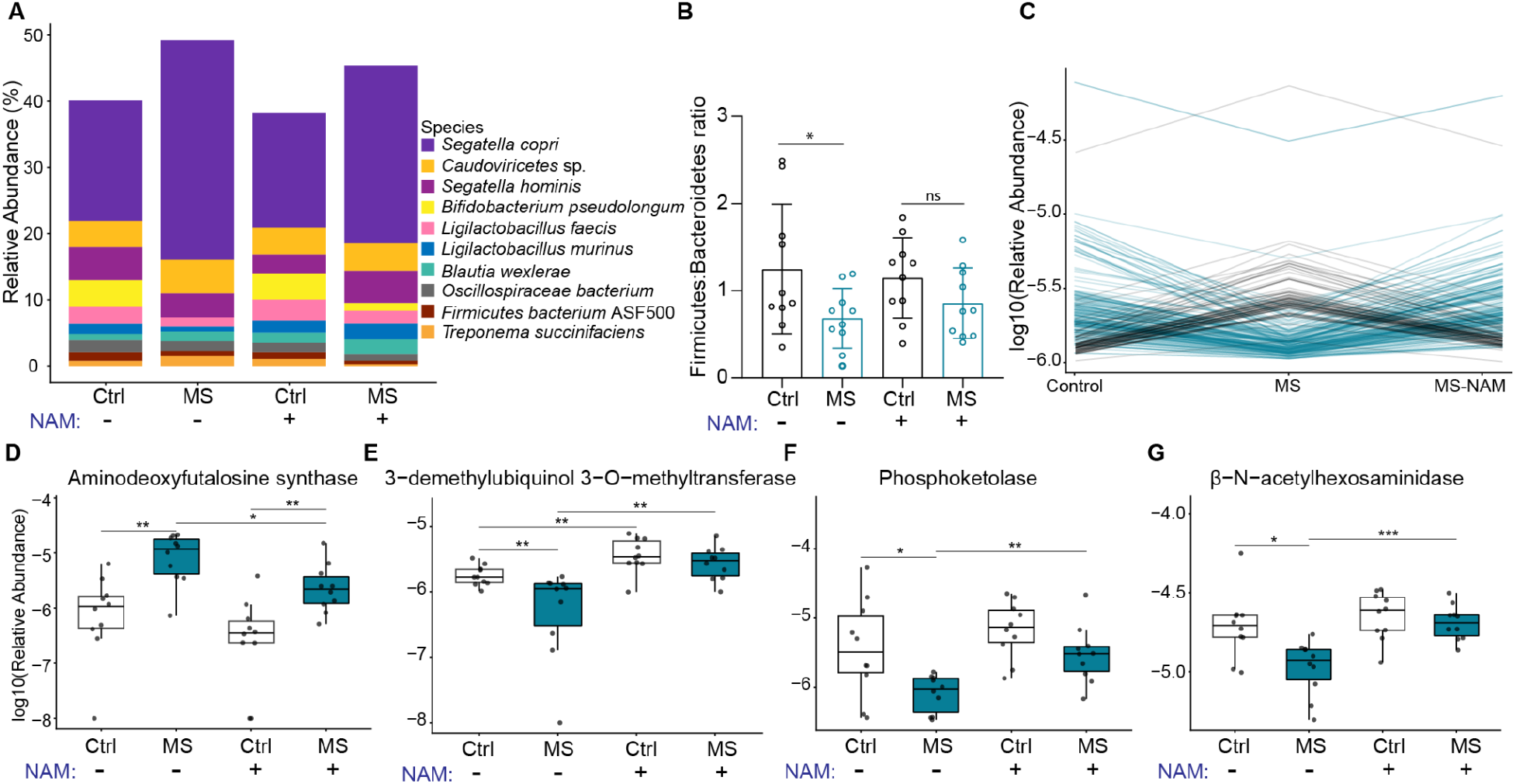
Early life stress and adult nicotinamide (NAM) supplementation impact gut microbiome composition and function. (A) Stacked bar plots showing the top ten most abundant species across groups represented on the x-axis. (B) Bar plots showing Firmicutes to Bacteroidetes ratios for different groups. Each data point represents an independent biological replicate used for shotgun sequencing. Error bars show standard deviation. (C) Line plot showing the log_10_(relative abundance) of COGs following the pattern of increase/decrease with MS and decrease/increase with NAM supplementation, represented by black or blue lines, respectively. Each line represents a different COG. (D-G) Boxplots showing the log_10_(relative abundance) for a few representative enzyme commission (EC) numbers. Horizontal lines represent the median. (For A-G: n=10 biological replicates; for B and D-G: statistics were performed using a Kruskal-Wallis test with a Benjamini-Hochberg post-hoc adjustment. ns: *p*-value > 0.05, *: *p*-value ≤ 0.05, **: *p*-value ≤ 0.01, and ***: *p*-value ≤ 0.001).

Next, to assess whether the premature aging effects of MS apply to the gut microbiome, we calculated the Firmicutes to Bacteroidetes (FB) ratios in different samples. This ratio is reported to increase from childhood to adulthood and then decrease during old age, likely due to inflammation and metabolic changes^25,26^. The FB ratio of maternally separated animals was significantly lower compared to healthy controls in middle age (Fig 2B). We observed a trend of increasing FB ratio upon nicotinamide supplementation in animals that underwent MS.

Further, we used HUMAnN3 to examine the differentially abundant clusters of orthologous genes (COGs) after MS and nicotinamide supplementation (Suppl. Fig. 2A-B) ^27^. We applied *p*-value (≤0.05) and fold change (≥ 2) cut-offs to filter COGs with significant changes in abundance. We observed ∼7% of the significantly altered COGs to show a supplementation-responsive trend (Fig. 2C). The nicotinamide supplementation increased the abundance of 176 COGs that were decreased after MS and decreased the abundance of 75 COGs increased in abundance after MS. This result indicates that nicotinamide might selectively promote taxa associated with potentially critical functions in the gut microbiomes of maternally separated rodents.

Gut bacteria contribute to host NAD metabolism through microbial nicotinamidase-mediated conversion of nicotinamide to nicotinic acid, which subsequently enters the Preiss-Handler pathway for NAD biosynthesis ^28^. In addition, many bacteria can synthesize NAD directly from nicotinamide through their own salvage pathways ^19^. Consequently, nicotinamide supplementation not only affects host metabolism but also alters the composition and metabolic activity of the gut microbiome ^29^. In our previous study, we observed restoration of energy homeostasis in maternally separated animals on nicotinamide supplementation ^16^. These observations prompted us to investigate whether early-life stress and nicotinamide supplementation reshape the bioenergetic profile of the gut microbiome also. Notably, the inflammatory milieu associated with MS can directly influence microbial bioenergetics and respiratory metabolism ^30^. We therefore examined COGs related to energy metabolism in maternally separated and nicotinamide-supplemented animals (Suppl. Fig. 2C-D).

In animals with a history of maternal separation, we observed an increase in the abundance of COGs related to hypoxic or anaerobic respiration including the nitrate reductase, cytochrome-*bd* oxidase, and menaquinone biosynthetic enzymes (Suppl. Fig. 2C). The elevation of bacterial taxa harboring nitrate reductase aligns with previous reports showing that maternal separation promotes gut inflammation ^31^. Nitrate is an alternate electron acceptor produced as a byproduct of the host immune response in inflamed niches ^30,32^. Interestingly, the menaquinone biosynthesis COGs which increase in abundance upon MS belong to the futalosine-dependent pathway which is canonically less prevalent in commensal microbes of healthy gut ^33^. Nicotinamide supplementation appears to alleviate many of these changes in COG abundances (Suppl. Fig. 2D). Thus, nicotinamide potentially promotes a shift in the functional state of the gut microbiome.

We further resolved the reaction-level differences caused by nicotinamide supplementation in maternally separated animals by mapping the reads from genome sequencing to corresponding Enzyme Commission numbers (ECNs) using HUMAnN3. Among the top 25 differentially abundant ECNs, we observed certain enzymes associated with energy metabolism, gut barrier physiology, and inflammation (Fig. 2D-G). Consistent with our COG analysis, the futalosine-dependent menaquinone biosynthesis enzyme, aminodeoxyfutalosine synthase, was significantly increased in maternally separated animals but decreased after nicotinamide supplementation. We found the ubiquinone biosynthesis enzyme 3-demethylubiquinol 3-O-methyltransferase to be decreased with MS and increased upon nicotinamide supplementation. Phosphoketolase, which is a key enzyme for the pentose-phosphate pathway (PPP), was also depleted with MS. Earlier, we had observed depletion of several glycolysis and PPP-associated COGs upon MS. We also found a decrease in □-N-acetylhexosaminidase, an enzyme involved in the hydrolysis of complex polysaccharides such as those present in mucin. This enzyme aids in the regulation of anti-inflammatory T-lymphocytes in the intestines, and the breakdown products of its catalytic reaction are critical for the maintenance of gut barrier integrity ^34,35^. We found that the abundance of □-N-acetylhexosaminidase is increased after nicotinamide supplementation, indicating potential protective effects on gut barrier permeability.

Together, our observations indicate that gut microbial dysbiosis induced by early-life stress persists well into middle age and is accompanied by broad alterations in microbial functional potential. Adult nicotinamide supplementation partially reverses these taxonomic and functional signatures. Although these findings do not establish causal mechanisms, they suggest that microbiome remodeling may contribute to the beneficial effects of nicotinamide observed in models of early-life stress. More broadly, our work motivates mechanistic studies to determine whether targeting host microbiome metabolic interactions can mitigate the long-term consequences of early-life adversity and inform future strategies for managing early-life stress-associated disorders.

## Supporting information

Detail of Cluster of Orthologous Genes

## Materials and Methods

Detailed in the supplementary file.

## Data availability

The 16S rRNA amplicon sequencing and whole genome shotgun sequencing data are available on NCBI Sequence Read Archive (16S: PRJNA1479155, WGS: PRJNA1480460).

## Code availability

All the pipelines used in this work are cited to the original authors.

## Acknowledgements

This work was funded by the DAE, India-Tata Institute of Fundamental Research Grant to A.A. (RTI4015) and V.A.V. (RTI4015, RTI4003). We acknowledge funding to VAV from the Sree Ramakrishna Paramahamsa Research Grant for Translational Biomedical Research (SreePVF/G/BS/19/1) and the J.C. Bose Fellowship to VAV (Grant no. JCB/2021/000014). P.R.C. acknowledges funding support from India Alliance (Department of Biotechnology, India and Wellcome Trust, UK) Early Career Fellowship (Reference No. IA/E/18/1/504310). Icons for some figures were created using our license at Biorender.com (ANAND, A. (2026) https://BioRender.com/8l9hfur). We thank TIFR animal facility staff for their help with animal experiments.

## Author contributions

Conceptualization: S.S. and A.A.; Supervision: V.A.V. and A.A.; Funding: A.A and V.A.V.; Methodology: S.S., P.R.C., Sh.S., V.A.V., and A.A.; Investigation: S.S., P.R.C., and N.P.; Visualization: S.S. and A.A.; Writing the original draft: S.S. and A.A.

