## Supplementary material for "Early-life stress-induced gut dysbiosis is ameliorated by nicotinamide treatment": Methods and supplementary figures

### Materials and methods:

#### Subjects

Sprague-Dawley (SD) rats (*Rattus norvegicus*) were bred in the Tata Institute of Fundamental Research (TIFR) animal facility. All animals were group housed and maintained on a 12:12 hour light:dark cycle (lights on at 07:00) with access to food and water *ad libitum*. Animal procedures were in accordance with the Committee for Control and Supervision of Experiments on Animals (CCSEA) guidelines and were approved by the Institutional Animal Ethics Committee (TIFR/IAEC/2024-1). Care was taken across all experiments to minimize animal suffering and restrict the number of animals used.

#### Maternal separation paradigm

Animals were subjected to maternal separation (MS) from postnatal day (PND)2 to PND14 as described previously[1,2]. Briefly, litters born to pregnant primiparous dams were assigned randomly to control or MS groups on PND1. All experimental litter sizes were maintained at 10 pups. Pups in the MS group were separated as a litter from their mothers for 3 hours daily (10:00 to 13:00) from PND2 to PND14. The dams from the MS group were first removed from their home cage to a novel cage prior to the removal of the litter. During this period, pups were placed in beakers with bedding and nesting material similar to their home cage and the beakers were placed on temperature controlled heating pads (Flamingo, India) to maintain euthermic conditions. Pups were returned to their home cage at the end of the separation period prior to reinstatement of the dam. Control litters and dams were left undisturbed in their home cage except for routine animal facility rearing. All the pups, from both control and MS groups, were weaned at PND28 and then housed in sex-matched groups of 3 to 4 animals per cage. Subsequent experiments were carried out in young-adult (2 month old) and middle-aged (13 month old) male rats from control and MS litters.

#### Oral nicotinamide supplementation

8-month old control and MS rats were subjected to nicotinamide (#72340, Sigma Aldrich, USA) administration (100mg/kg/day) in drinking water for a 5-month period as described previously [1]. Concomitantly, a cohort of control and MS animals continued receiving regular drinking water.

The bottles were changed every day and animal weights were monitored every 3-4 weeks. Middle-aged control and MS animals with or without nicotinamide administration were used for the collection of fecal samples for further investigations.

**Collection of fecal samples for sequencing**

At young adult and middle-aged time points (2 months old and 13 months old, respectively), rats from the control, MS, and nicotinamide supplemented groups were used for collecting fecal samples. All sample collections were performed at approximately similar time of the day (10:00-10:30). The animal of interest was separated into a novel cage and allowed to defecate, following which it was returned to its respective home cage. Fresh fecal pellets were immediately collected in a sterile cryovial and stored on ice until the samples were transferred to -80°C as soon as possible. The animals were then returned to their respective home cages. Fecal samples were processed for isolating DNA and further processing.

**DNA extraction**

150 mg of fecal sample was taken for DNA extraction. Lysis was performed using the Qiagen TissueLyser, followed by DNA isolation according to the manufacturer's protocol using the QIAamp PowerFecal Pro DNA Kit (#51804, Qiagen GmbH, Germany). The DNA samples were quantified on NanoDrop One and Qubit using water and standard as controls respectively.

**16S rDNA V3-V4 metagenomic library preparation**

Libraries were constructed in alignment with the 16S metagenomic library preparation protocol from Illumina Inc. The gene-specific sequences used in this protocol targeted the 16S V3 and V4 region. Illumina adapter overhang nucleotide sequences were added to the gene-specific sequences. The full length primer sequences targeting the V3-V4 region were as follows:

| Name | Sequence |
|---|---|
| 16S Amplicon PCR Forward Primer | 5'TCGTCGGCAGCGTCAGATGTGTATAAGAGACAGCCTACGGGN GGCWGCAG3' |

| | |
|---|---|
| 16S Amplicon PCR Reverse Primer | 5'GTCTCGTGGGCTCGGAGATGTGTATAAGAGACAGGACTACHVGGGTATCTAATCC3’ |

Briefly, 12.5 ng DNA was subjected to 16S V3-V4 PCR using the respective primers, the PCR products were bead purified and subjected to another round of PCR with dual indices and adapters to generate the respective libraries. The cleaned libraries were quantitated on Qubit fluorometer and appropriate dilutions loaded on a D1000 screen tape to determine the size range of the fragments and the average library size. All the libraries were taken forward for sequencing using the Illumina NextSeq 2000 platform for 2 x 300 bp pair-ended sequencing.

**Shotgun whole genome sequencing library preparation**

Libraries were constructed in alignment with whole genome sequencing recommendations of Twist Library Preparation EF Kit 2.0 (#101058). Briefly, 100 ng of DNA was used for fragmentation and end repair, followed by universal adapter ligation, removal of excess adapters using Twist purification beads, PCR enrichment of adapter-ligated DNA and clean-up of library products as instructed by the manufacturer. The cleaned libraries were quantitated on Qubit fluorometer and appropriate dilutions loaded on Labchip to determine the size range of the fragments and the average library size. All the libraries were taken forward for sequencing using the Illumina NextSeq 2000 platform for 2 x 150 bp pair-ended sequencing.

**Data processing for 16S rRNA amplicon sequencing**

The raw reads were trimmed & filtered using Fastp v1.0.1 [3] to remove adaptors, low quality reads (Q<30). Data was imported into QIIME2 [4] and the quality of the paired-end reads was inspected. The sequence denoising and primer removal was performed with the denoise-paired method of the DADA2 plugin of QIIME2. The denoising was done using a 17 bp left trim and 21 bp right trim for primer removal, along with truncation to 290 bp and 270 bp for the forward and reverse reads, respectively, to remove low-quality bases at the end of the reads.

A total of 13,703 ASVs (amplicon sequence variants) were detected. The mean length was found to be 408.51 bp. The database used for further processing contained backbone 16S rRNA sequences from Greengenes2 [5]. The feature-table and representative sequences from the denoising steps were used as input along with the downloaded reference database. The "non-v4-16s" plugin of Greengenes2 was used to perform closed reference OTU picking, using q2-vsearch, against the full length 16S sequences in Greengenes2. This led to a significant reduction in the number of ASVs (a total of 1527 ASVs with a mean length of 408.75bp). The data were imported in tabular format to create a rarefied phyloseq [6] object in R for downstream data analysis and visualization.

**Data processing for shotgun sequencing**

Fastp v1.0.1 [3] was used for quality trimming and FastQC v0.12.1  for quality assessment. The trimming parameters applied were a Q score ≥ 30 and a minimum read length of 50. To eliminate host-derived sequences for downstream metagenomic analyses, the processed reads were aligned to the *Rattus norvegicus* (GRCr8: GCF_036323735.1) reference host genome using Bowtie2 v2.4.4 [7]. Reads that aligned to the host genome were removed and only the host-filtered reads were retained. These host-filtered reads formed the input for metagenomic profiling and taxonomic classification.

The host-filtered reads were classified using Kraken2 v 2.17.1 [8] against the Core_NT (2025) database. The relative abundance was calculated at each taxonomic level.

**Alpha and beta diversity measurement**

The statistical comparison for Chao1, Shannon, and Simpson diversities was carried out using the Wilcoxon rank-sum test. All the plots and comparisons for alpha diversity, Bray-Curtis PCoA and Jaccard NMDS were carried out using the phyloseq R package.

**Taxonomic abundance measurement**

For 16S rRNA amplicon sequencing data, the merged ASV table was used to calculate the relative taxonomic abundance. From here, the taxa were separated and the ASVs belonging to the same taxonomic classification were merged to get the final abundance table, which is then used for the relative abundance calculation. Taxonomic lineages not included in the top 10 were collapsed as "Others". A similar approach was used for the shotgun sequencing data, except that BIOM files generated as the output of Kraken2 were used to perform taxonomic classification.

**Read-based functional profiling**

Functional profiling was performed using HUMAnN3 v3.9[9]. The abundances in reads per kilobase (RPK) of gene families obtained from HUMAnN analysis were renormalized to relative abundance. The gene families were then grouped into Gene Ontology (GO) terms, EC categories, EggNOG (including COG) terms and MetaCyc reactions using HUMAnN3 Utility scripts.

**Statistical analysis**

Data and statistical analysis was performed in R v4.4.1 using packages ggplot2 and VEGAN [10–12]. To calculate significance between two groups a Wilcoxon rank-sum test was performed. To calculate significance between groups a Kruskal-Wallis test with false discovery rate (FDR) correction, using the Benjamini and Hochberg method, was applied. For alpha and beta diversity analyses sequences were rarified to equal sequencing depth. Significance level was set to $p \leq 0.05$. For COG changes, significance extraction was performed by setting a prevalence filter of 70%, a $\log_2$(fold change) cut-off of 1 or -1, and a p-value cut off of 0.01 (Supplementary sheet 1).

### Supplementary Figures

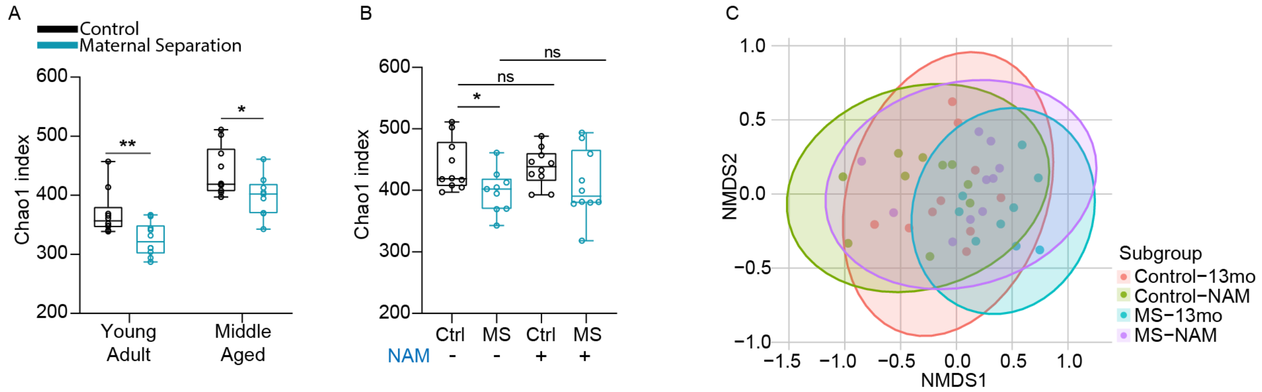


**Supplementary Figure 1:** Gut microbiome diversity alterations associated with maternal separation and nicotinamide (NAM) supplementation. (A) Box plot representing changes in Chao1 index in gut microbiomes from control and MS animals at young adulthood and middle age timepoints. (B) Box plot representing changes in Chao1 index in gut microbiomes from control and MS animals at middle age with NAM supplementation. In A&B: n=9-10, box plots extend from 25th to 75th percentiles; whiskers represent minimum and maximum data points. Statistics were performed using the Wilcoxon rank-sum test with Bonferroni post-hoc correction. ns: p-value > 0.05, *: p-value ≤ 0.05, and **: p-value ≤ 0.01. (C) A Jaccard index-based non-metric multidimensional scaling (NMDS) plot for control and MS animals at middle age with and without NAM supplementation.

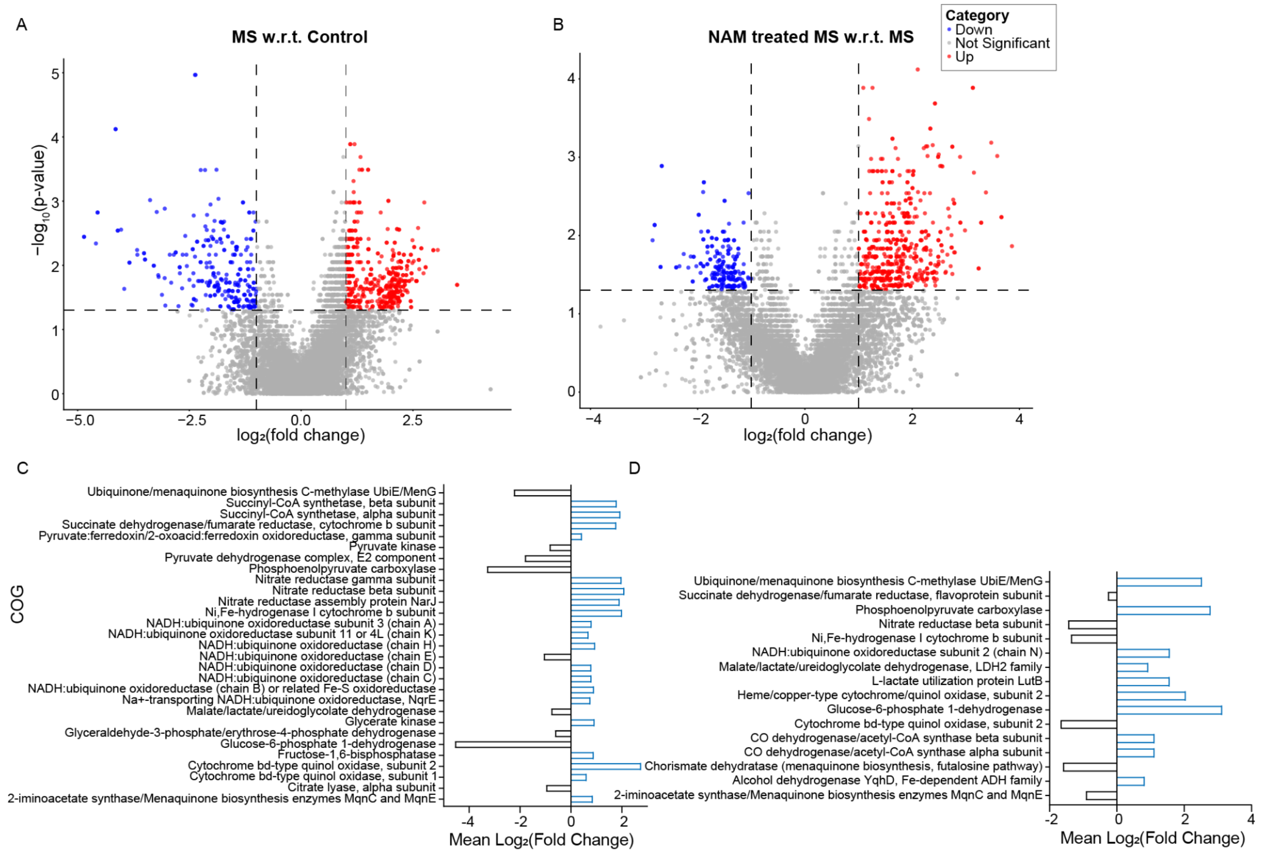

**Supplementary Figure 2:** Changes in clusters of orthologous genes (COGs) due to maternal separation and nicotinamide (NAM) supplementation. (A) Volcano plot depicting relative abundance changes for COGs in maternally separated animals compared to controls. (B) Volcano plot depicting relative abundance changes for COGs in maternally separated animals with and without nicotinamide supplementation. In A&B: fold change cut-off was set at ≥2 and *p*-value cut-off was set at ≤0.05. Significance was assessed using a Wilcoxon rank sum test with the Benjamini-Hochberg post-hoc correction. (C) Bar plot showing the change in mean $\log_2$(fold change) for energy metabolism-related COGs in maternally separated animals compared to controls. (D) Bar plot showing the change in mean $\log_2$(fold change) for energy metabolism-related COGs in maternally separated animals with and without nicotinamide supplementation.
